# Adaptive Multi-Objective Control Explains How Humans Make Lateral Maneuvers While Walking

**DOI:** 10.1101/2022.03.21.485079

**Authors:** David M. Desmet, Joseph P. Cusumano, Jonathan B. Dingwell

## Abstract

To successfully traverse their environment, humans often perform maneuvers to achieve desired task goals while simultaneously maintaining balance. Humans accomplish these tasks primarily by modulating their foot placements. As humans are more unstable laterally, we must better understand how humans modulate lateral foot placement. We previously developed a theoretical framework and corresponding computational models to describe how humans regulate lateral stepping during straight-ahead continuous walking. We identified goal functions for step width and lateral body position that define the walking task and determine the set of all possible task solutions as Goal Equivalent Manifolds (GEMs). Here, we used this framework to determine if humans can regulate lateral stepping during *non*-steady-state lateral maneuvers by minimizing errors consistent with these goal functions. Twenty young healthy adults each performed four lateral lane-change maneuvers in a virtual reality environment. Extending our general lateral stepping regulation framework, we first re-examined the requirements of such transient walking tasks. Doing so yielded new theoretical predictions regarding how steps during any such maneuver should be regulated to minimize error costs, consistent with the goals required at each step and with how these costs are adapted at each step during the maneuver. Humans performed the experimental lateral maneuvers in a manner consistent with our theoretical predictions. Furthermore, their stepping behavior was well modeled by allowing the parameters of our previous lateral stepping models to adapt from step to step. To our knowledge, our results are the first to demonstrate humans might use evolving cost landscapes in real time to perform such an adaptive motor task and, furthermore, that such adaptation can occur quickly – over only one step. Thus, the predictive capabilities of our general stepping regulation framework extend to a much greater range of walking tasks beyond just normal, straight-ahead walking.

**AUTHOR SUMMARY:** When we walk in the real world, we rarely walk continuously in a straight line. Indeed, we regularly have to perform other tasks like stepping aside to avoid an obstacle in our path (either fixed or moving, like another person coming towards us). While we have to be highly maneuverable to accomplish such tasks, we must also maintain balance to avoid falling while doing so. This is challenging because walking humans are inherently more unstable side-to-side. Sideways falls are particularly dangerous for older adults as they can lead to hip fractures. Here, we establish a theoretical basis for how people might accomplish such maneuvers. We show that humans execute a simple lateral lane-change maneuver consistent with our theoretical predictions. Importantly, our simulations show they can do so by adapting at each step the same step-to-step regulation strategies they use to walk straight ahead. Moreover, these same control processes also explain how humans trade-off side-to-side stability to gain the maneuverability they need to perform such lateral maneuvers.

## INTRODUCTION

To successfully traverse our environment, we humans must adapt to a wide variety of environment contexts and changing task goals [1] (e.g., Fig. 1A), all while maintaining balance. Indeed, humans readily avoid obstacles [2] and/or other humans [3], step to targets [4–6], move laterally [7–9], navigate complex terrain [10–12], and respond to destabilizing perturbations [13, 14]. To do so requires a high degree of maneuverability [7] and, potentially, the ability to trade-off stability for maneuverability [8, 9]. While maneuverability has been widely studied in animal locomotion [15, 16] and robotics [16, 17], far less is known about how humans accomplish such tasks.

**Figure 1.**
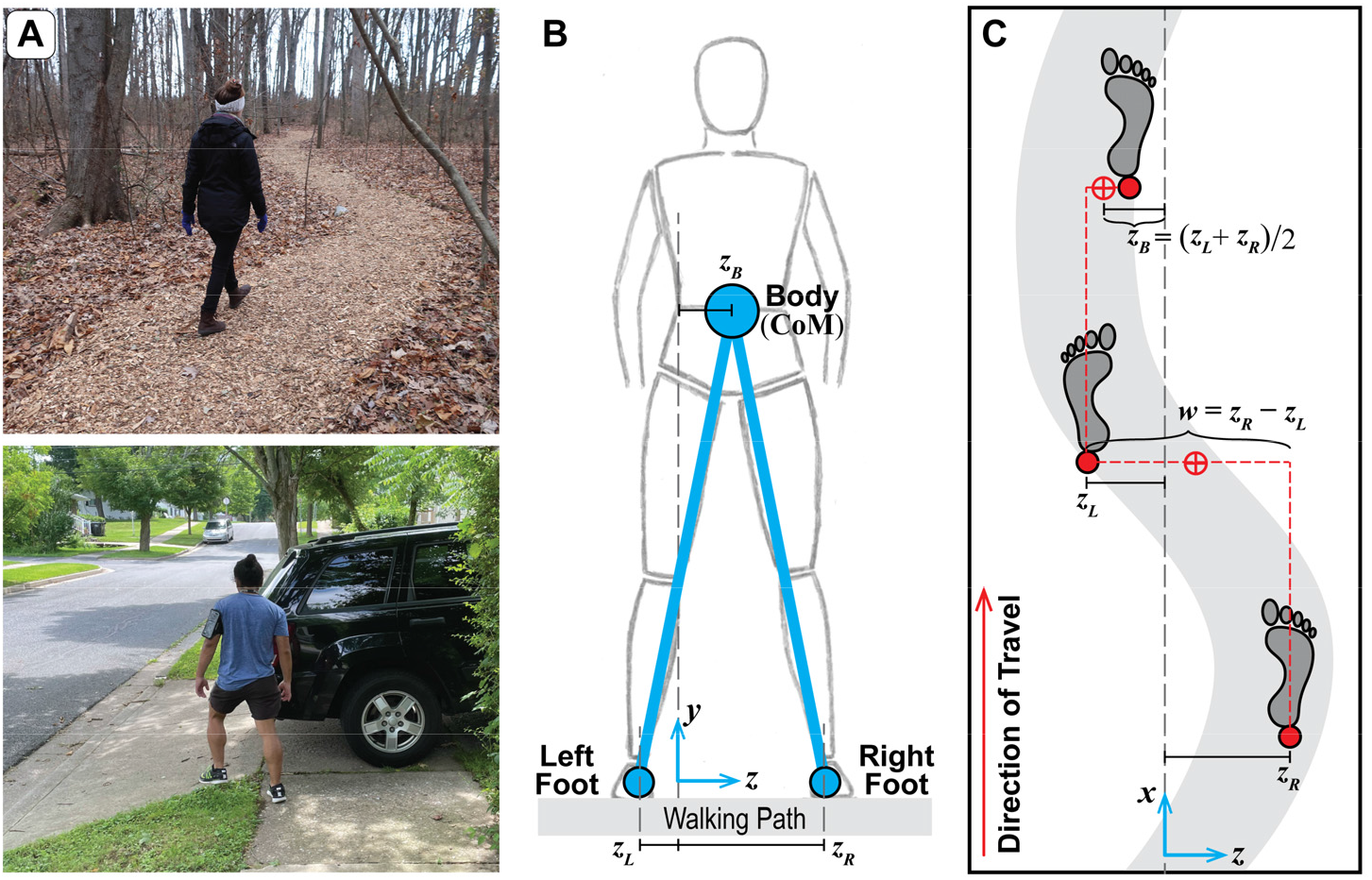
Defining Relevant Lateral Stepping Variables. A) Examples of people walking in common contexts that require adaptability, including walking on a winding path and avoiding an obstacle. B-C) Configuration of bipedal walking in both the frontal (B) and horizontal (C) planes. Coordinates are defined in a Cartesian system with {*x,y,z*} axes shown in (B) and (C) and the origin defined as the geometric center of the available walking surface. The lateral positions of the left and right feet {*z_L_, z_R_*} are used to derive the primary lateral stepping variables that are regulated from step-to-step: step width (*w*=*z_R_* – *z_L_*) and lateral body position (*z_B_* = ½(*z_L_* + *z_R_*)), which reflects a once-per-step proxy for the lateral position of the center-of-mass (CoM) ([1]; see Supplement).

Humans accomplish these sorts of walking maneuvers primarily by modulating their foot placements. Appropriate foot placement at each step can redirect center-of-mass accelerations, which enables humans to maintain balance while walking straight ahead [18–20]. This is especially important in the lateral direction where humans are thought to be more unstable [21–23]. Indeed, laterally-directed falls are particularly injurious in older adults [24, 25]. However, the express purpose of any lateral maneuver is to specifically interrupt ongoing straight-ahead walking to achieve some particular walking task goal that requires a walker to alter their foot placements [8, 26]. Thus, it is important to better understand how humans adapt their lateral foot placements to maintain balance while simultaneously executing such lateral maneuvers.

Human walking exhibits considerable variability [27, 28], redundancy (i.e., the body has more mechanical degrees of freedom than necessary to perform any movement) [29], and equifinality (i.e., humans can perform most tasks an infinite number of ways) [30, 31]. Computational models are necessary to fully understand how humans perform accurate, goal-directed walking movements in the context of these challenges. Most prior models of human walking (e.g., [21, 32, 33]) address within-step “control,” defined here as the processes that drive the dynamics of each step to remain viable (i.e., to prevent falling) [34]. While ensuring viability is indeed necessary to walk [35, 36], it is not sufficient: humans engage in *purposeful* walking with a destination to reach and/or other tasks to achieve. Control models that “just walk” (i.e., remain viable) do not address how humans achieve such goal-directed walking.

We posit that locomotion is functionally hierarchical, consisting of both within-step control and step-to-step regulation. To compliment within-step control models, we developed a motor regulation framework to determine how humans adjust successive stepping movements [1, 30, 34, 35]. For a given walking task (e.g., Fig. 1A), we use this framework to propose goal functions that theoretically define the task and, accounting for equifinality, determine the set of all possible task solutions as Goal Equivalent Manifolds (GEMs) [37–40]. The goal functions are incorporated into task-level costs, which we then use in a stochastic optimal control formalism [41] to generate relatively simple, phenomenological models of step-to-step motor regulation (i.e., “motor regulation templates” [1, 30, 31]). For any proposed goal function, these templates predict how humans would attempt to drive that goal function to zero at the next step, thus achieving perfect task performance on average. For an appropriate choice of goal functions, these models successfully replicate human step-to-step dynamics in both the fore-aft [30] and lateral [1] directions. In the lateral direction in particular, multi-objective regulation of primarily step width (*w*) and secondarily lateral position (*z_B_*) captures human step-to-step dynamics during continuous straight-ahead walking (Fig. 1B-C) [1, 13, 42]. Whether we construct explicit models or not, this overall theory and its hierarchical control/regulation schema provides a powerful framework from which to interpret experimental results [13, 42–46].

However, humans rarely perform long bouts of straight-ahead, continuous walking [47, 48]. Instead, humans must frequently perform various kinds of locomotor maneuvers when walking in the real world (e.g., Fig. 1A) [7–9]. Indeed, older adults often fall during such maneuvers because they incorrectly transfer their body weight or they trip [49]. Because both such causes can be prevented with appropriate foot placement [18, 20, 22, 50], it is necessary to better understand how humans regulate foot placement as they execute typical lateral maneuvers. Here, we aimed to determine how humans regulate their stepping during a simple lateral maneuver: namely, a single lateral lane-change transition between periods of straight-ahead walking.

It is not clear *a priori* that our previous lateral stepping regulation framework can also emulate human stepping during lateral maneuvers. This framework was originally developed to model straight-ahead steady-state walking, under assumptions that step-to-step adjustments can be made without changing the fundamental structure of within-step control, and that deviations from perfect performance are small. These assumptions motivated us to select low-dimensional, single-step, linear regulators to model these step-to-step error-correcting processes [1, 30, 31]. However, walking tasks requiring substantial *non*-steady-state lateral maneuvers would seem to violate these assumptions. Lateral maneuvers that introduce large deviations from steady-state might, for example, require changes to the within-step control structure, or induce substantial nonlinearity. Lateral maneuvers might also require substantially greater impulses to execute, which may necessitate additional compensatory motor and/or kinematic contributions and could result in the need for higher model dimensionality. Furthermore, humans plan maneuvers more than one step in advance in some contexts [4, 5], which suggests the possible need for models that depend on more than one prior step. Thus, for any of these reasons, one might reasonably posit that humans regulate stepping using entirely different control schemes when executing lateral maneuvers far from steady state.

However, here we extend our previous theoretical framework to demonstrate how humans might *adapt* how they regulate lateral stepping to execute non-steady-state maneuvers without having to resort to some entirely different scheme. We first apply concepts from the general theory in a manner that satisfies the requirements of a non-steady-state lane change maneuver. Specifically, we allow the task goals, task-level costs, and the weighting between competing costs to adapt from each step to the next. We thus derive a model that yields explicit, empirically-testable predictions about how the variability observed during such lateral maneuver tasks will be structured from step to step. We then test these predictions against human experimental data. Lastly, we used the models with these additional adaptive hierarchical elements to simulate how such lateral maneuvers are achieved over a sequence of steps. In doing so, we demonstrate how humans modulate their stepping to trade-off lateral stability for maneuverability to accomplish the lane change task. Thus, our hierarchical motor regulation framework can be applied to a far wider range of walking tasks beyond just straight-ahead, steady-state walking.

## RESULTS

### Lateral Maneuvers in Humans

We analyzed data from a prior study conducted in our laboratory [51]. Young healthy participants walked in a virtual reality environment. They walked on one of two parallel paths, the centers of which were 0.6m apart, projected onto a 1.2m wide motorized treadmill (Fig. 2A). Following an audible cue, they executed a lateral maneuver from the path they were walking on over to the adjacent path. Importantly, participants were given *no* explicit instructions about how to execute this maneuver. We analyzed seventy-nine total maneuvers from twenty participants.

**Figure 2.**
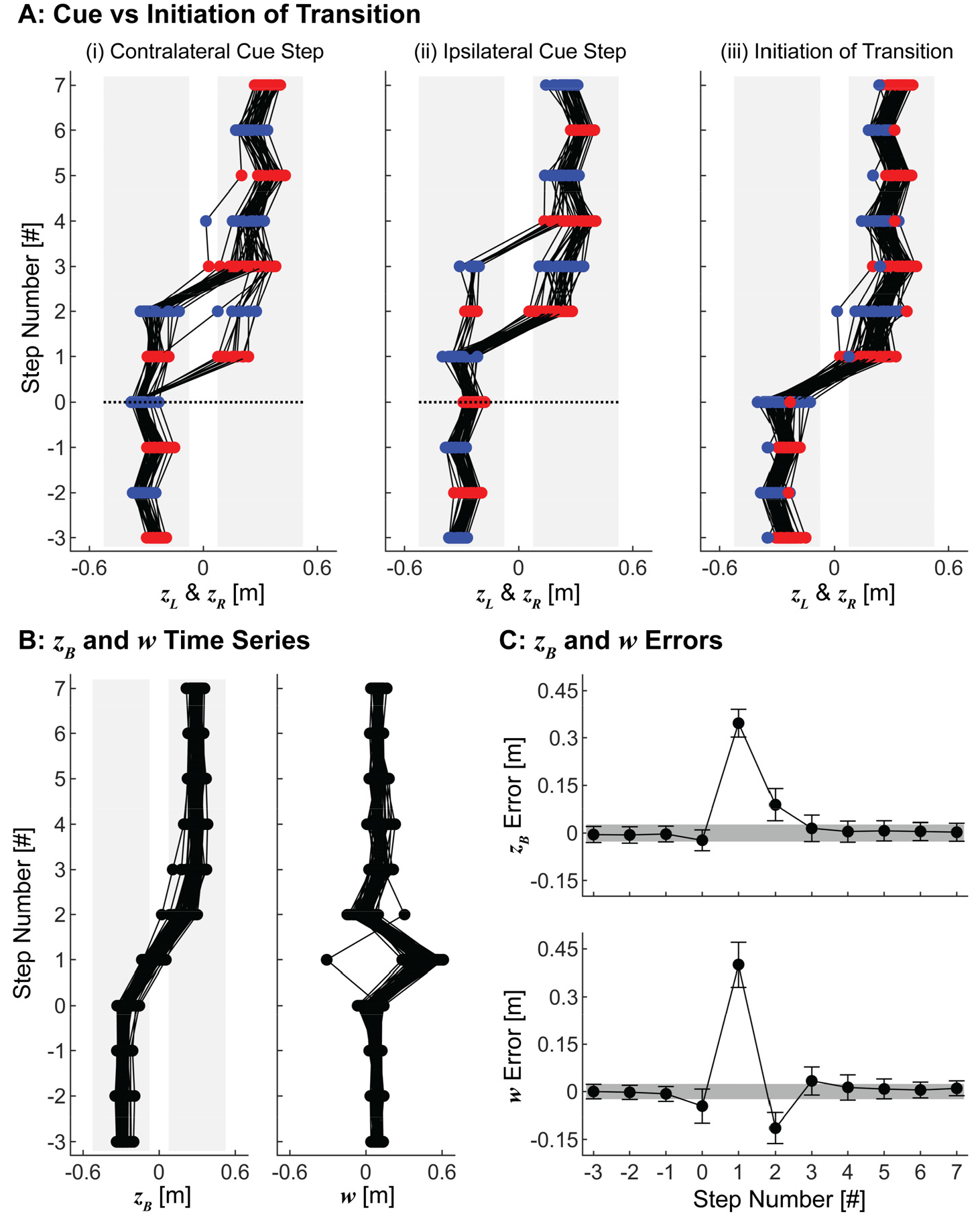
Experimental Stepping Time Series and Errors. A) Time series of left (*z_L_*; blue) and right (*z_R_* red) foot placements for (i) 38 transitions when the cue was given on a contralateral step relative to the direction of transition, (ii) 41 transitions when the cue was given on an ipsilateral step relative to the direction of transition, and (iii) all 79 transitions with respect to the initiation of the transition, defined as the last step taken on the original path. In (i)-(ii), the black dotted lines at step 0 indicate the onset of the audible cue. All transitions are plotted to appear from left to right. B) Time series of lateral position (*z_B_*) and step width (*w*) for all transitions with respect to the initiation of the transition, with all transitions plotted to appear from left to right. C) Errors with respect to the stepping goals, [*z_B_*\*, *w*\*]. For steps in the interval [−3, 0], *z_B_*\* was defined as the experimental mean *z_B_* during steady state walking before the transition. For steps in the interval [1, 7], *z_B_*\* was defined as the experimental mean *z_B_* during steady state walking after the transition. For all steps analyzed, *w*\* was defined as the experimental mean *w* during steady state walking. Error bars indicate experimental standard deviations at each step. Gray shaded regions indicate the mean standard deviation (± 1) from all steady state walking steps.

Participants variably took 0-to-3 steps between the cue and initiating the maneuver (Fig. 2A[i-ii]). However, once initiated, they performed each maneuver consistently (Fig. 2A[iii]-B). Participants completed nearly all maneuvers with an ipsilateral transition step that involved a single, large step to reach the new path. One participant, however, completed one of their maneuvers by taking a large cross-over step. Additionally, one maneuver from another participant required three steps to reach the new path. These two non-conforming maneuvers demonstrate that participants had many options: the task itself did not require them to complete the maneuver in any specific way.

Participants completed nearly all lateral maneuvers in 4 non-steady-state steps. Participants first took a “*preparatory*” step (step 0) prior to the transition, during which they slightly narrowed their step width and incrementally moved towards the new path. They then took a large “*transition*” step (step 1) to cover most of the transition distance. Participants took a subsequent “*recovery*” step (step 2) that again exhibited a slightly narrowed step width. Participants then reached their final new goal position (step 3) and returned to steady-state walking on their new path.

### Steady-State Stepping Regulation Cannot Replicate Lateral Transitions

We first used our previous multi-objective model of lateral stepping regulation to test whether humans could regulate stepping during lateral maneuvers using a constant regulation strategy like that used for continuous, straight-ahead walking [1]. This model selects each new foot placement (*z_L_* or *z_R_*) as a weighted average of independent predictions that minimize errors with respect to either a constant step width (*w*\*) or lateral position (*z_B_*\*) goal, consistent with multi-objective stochastic optimization of error costs with respect to these two quantities (see Methods). For this model, the relative proportion of step width to lateral position regulation was defined by *ρ*, where *ρ* = 0 indicates 100% *z_B_* control (and hence, 100% weight on the *z_B_* cost) and *ρ* = 1 indicates 100%*w* control (100% weight on the *w* cost) [1]. This stepping regulation model reproduced the key features of human stepping dynamics during continuous, straight-ahead walking for constant values in the approximate range 0.89 ≤ *ρ* ≤ 0.97 [1].

We assessed whether this model, with any constant value of *ρ*, could emulate the *non*-steady-state stepping dynamics experimentally observed during the lateral maneuver task. We found that it was not capable of doing so. The model did emulate steady-state stepping dynamics (Fig. 3C; Step −3) for constant values in the range of approximately 0.83 ≤ *ρ* ≤ 0.92, consistent with previous findings [1]. However, *no* simulations over this range of *ρ* completed the lateral maneuver as observed in the experiment (Fig. 3A-B; red). Furthermore, while model simulations that weighted position and step width regulation similarly (i.e., *ρ* ≈ 0.5) approximately emulated average experimentally observed stepping time series and errors (Fig. 3A-B; blue), they failed to emulate the stepping variability humans exhibited during either steady-state (Step −3) or transition (Step 1) steps (Fig. 3C). Indeed, no single constant value of *ρ* emulated both steady-state and transition stepping dynamics (Fig. 3C).

**Figure 3.**
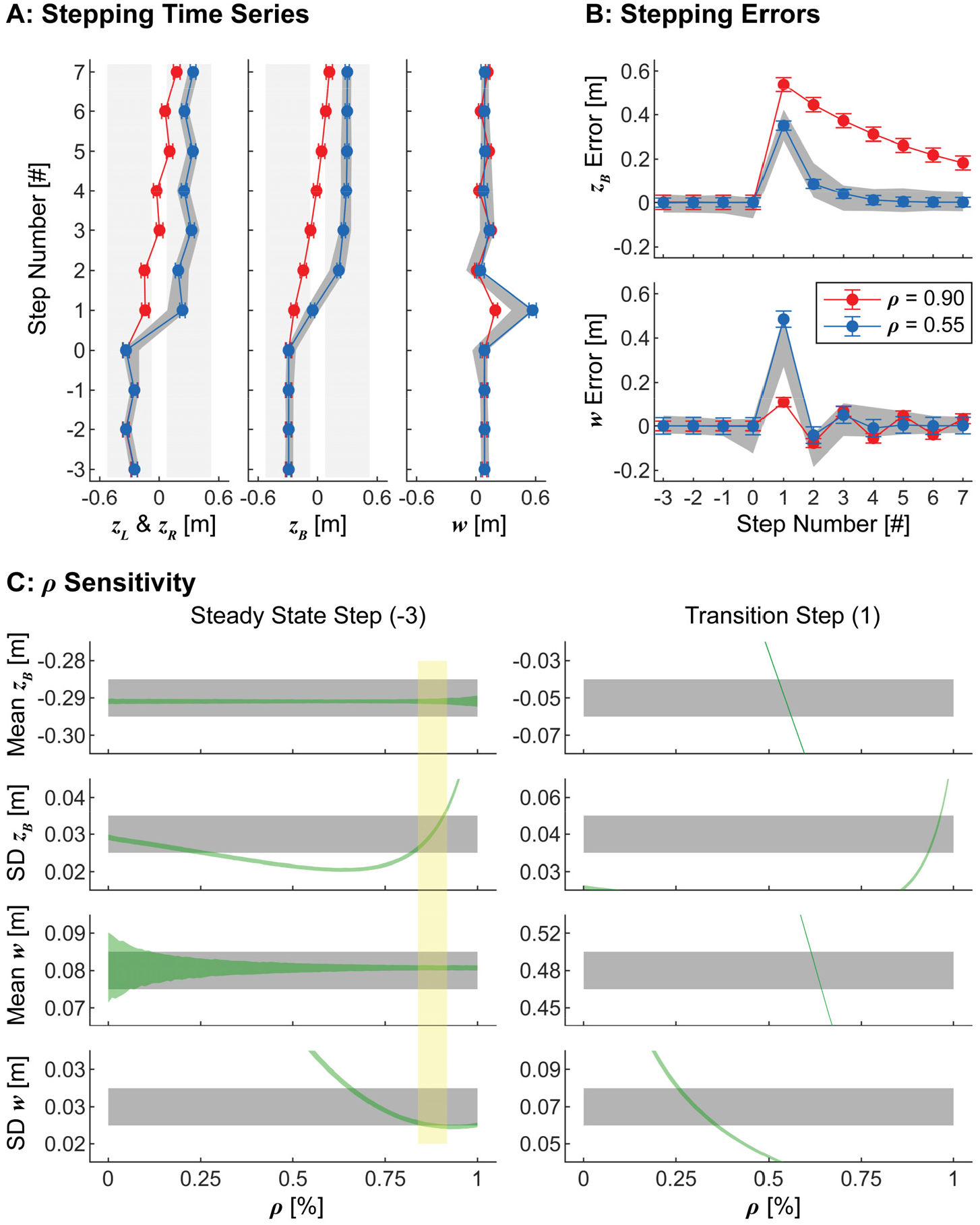
Constant Parameter Model Results. A) Simulated stepping time series (mean ± SD) of 1000 lateral transitions using the original, constant parameter model with *ρ* = 0.90 (red) and *ρ* = 0.55 (blue). For steps in the interval [−3, 0], we set *z_B_*\* as the experimental mean *z_B_* during steady state walking before the transition. For steps in the interval [1, 7], we set *z_B_*\* as the experimental mean *z_B_* during steady state walking after the transition. For all steps, *w*\* was defined as the experimental mean *w* during steady state walking. B) Stepping errors (mean ± SD) at each step relative to the stepping goals, [*z_B_*\*, *w*\*], for the same data as in (A). For both (A) and (B), gray bands indicate the middle 90% range from experimental data. C) Means and standard deviations of both regulated variables (*z_B_* and *w*) during a steady state step (Step −3; left) and during the transition step (Step 1; right) for all values of 0 ≤ *ρ* ≤ 1. Gray bands indicate 95% confidence intervals from the experimental data computed using bootstrapping. Green bands indicate ±1 standard deviation from 1000 model simulations at each value of *ρ*. Model simulations over the approximate range of 0.83 ≤*ρ* ≤ 0.92 fell within the experimental ranges for all variables for steady-state walking (Step −3; Left), as indicated by the region highlighted in yellow. However, no such range captured the experimental data during the transition step (Step 1; Right).

### Re-Thinking Stepping Regulation for Non-Steady-State Tasks

Any biped (human, animal, robot, etc.) must enact step width and/or lateral position regulation via left and right foot placement (Fig. 1C). The stepping goals, [*z_B_*\*, *w*\*] that guide *steady-state* walking individually form diagonal and orthogonal Goal Equivalent Manifolds (GEMs) when plotted in the [*z_L_, z_R_*] plane (Fig. 4A) [1]. The intersection of these GEMs represents the multi-objective goal to maintain both *z_B_*\* and *w*\* and therefore defines the foot placement goal, [*z_L_*\*, *z_R_*\*], for the task. Along any GEM, deviations tangent to the GEM are “goal equivalent” because they do *not* introduce errors with respect to the goal. Conversely, deviations perpendicular to the GEM are “goal relevant” because they *do* introduce such errors [30]. Humans typically exhibit greater variability along GEMs they exploit [37, 41, 52]. Here, because the *z_B_*\* and *w*\* GEMs are orthogonal, goal equivalent deviations with respect to either GEM are goal relevant with respect to the other [1]. Hence, the ratio of the *δ_zB_* and *δ_w_* deviations with respect to both the *z_B_*\* and *w*\* GEMs theoretically reflects the relative weighting of *z_B_* and *w* regulation. During steady-state walking, the distribution of human steps is strongly *anisotropic* (Fig. 4A): steps are strongly aligned along the *w*\* GEM (such that *δ_z_B*\*/*δ_w_*>> 1) because humans heavily weight regulating step width over position [1].

**Figure 4.**
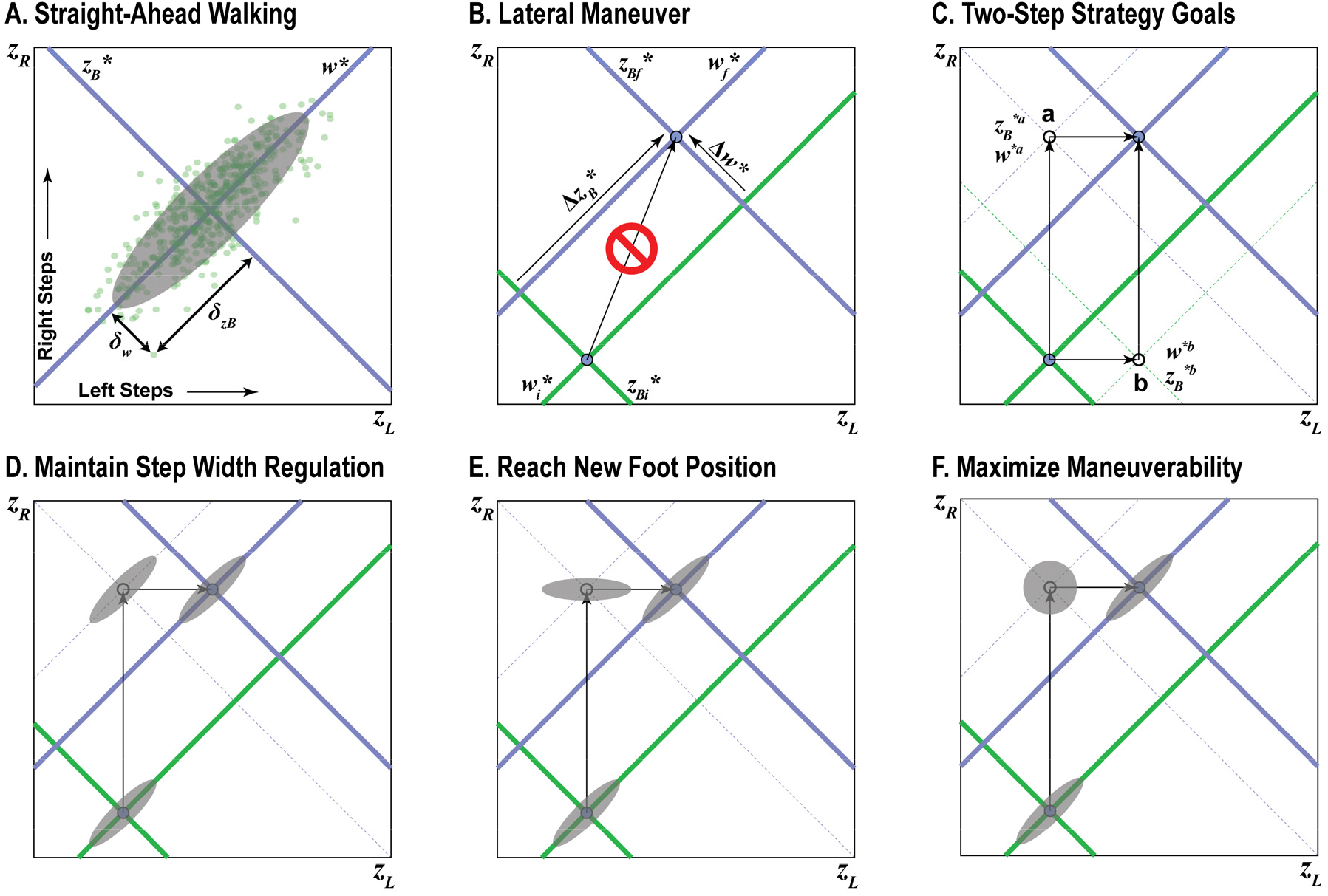
Conceptual Depiction of Task Performance. A) When viewed in the [*z_L_, z_R_*] plane, goals to maintain constant position (*z_B_*\*) or step width (*w*\*) each form linear Goal Equivalent Manifolds (GEMs) that are diagonal to the *z_L_* and *z_R_* axes and orthogonal to each other. Deviations (*δz_B_* and *δw*) with respect to both the *z_B_*\* and *w*\* GEMs characterize the stepping distribution at a given step and reflect the relative weighting of *z_B_* and *w* regulation. In steady-state walking, humans strongly prioritize regulating *w* over *z_B_*, producing stepping distributions strongly aligned to the *w*\* GEM. B) Any maneuver would then involve a substantial change from some initial (green) to some new final (blue) stepping goals that will displace these GEMs diagonally in the [*z_L_, z_R_*] plane, such as the theoretical rightward shift in *z_B_*\* (Δ*z_B_*\*) and increase in *w*\* (Δ*w*\*) depicted here. To accomplish such a maneuver requires changing both *z_L_* and *z_R_*. This cannot be achieved in any single step. C) At least two consecutive steps (either ‘a’: *z_R_* → *z_L_*, or ‘b’: *z_L_* → *z_R_*) at minimum are required to execute a lateral maneuver (Δ*z_B_*\* and/or Δ*w*\*). Each possible intermediate step (‘a’ or ‘b’) has its own distinct stepping goals. D-F) For any given intermediate step, numerous feasible strategies to execute the maneuver are theoretically possible. D) One such strategy might be to maintain strong prioritization of *w* over *z_B_* regulation (i.e., as in A) at the intermediate step. Enacting this strategy would produce a stepping distribution at the intermediate step that would remain strongly aligned to the new constant step width GEM (*w*^*a^) at that intermediate step. E) Another feasible strategy might be to simply put the first (here, right) foot at its new desired location (*z_R_*). This would produce a stepping distribution at the intermediate step that would be strongly aligned to that new desired location (here, *z_R_*) for that step. F) A third feasible strategy might be to maximize maneuverability. Here, foot placement at the intermediate step should be as accurate as possible. This would produce an approximately isotropic (i.e., circular) stepping distribution at the intermediate step.

When viewed in this manner, it then becomes evident that nearly any substantive change in either stepping goal (i.e., *Δz_B_*\* and/or Δ*w\**) will induce a diagonal shift of the corresponding GEM(s) in the [*z_L_, z_R_*] plane (Fig. 4B). This will necessitate corresponding changes in both left *and* right foot placement, which therefore cannot be accomplished in a single step (Fig. 4B). At minimum, two consecutive steps are required to execute nearly any maneuver involving some Δ*z_B_*\* and/or Δ*w*\*. The first step must be taken by either the left or right foot to either of two possible intermediate foot placement goals in the [*z_L_, z_R_*] plane (Fig. 4C). Indeed, we can derive exact stepping goals for this intermediate step algebraically (see Methods). However, equifinality exists in both the number and placements of steps that can be used to accomplish any Δ*z_B_*\* and/or Δ*w*\* maneuver. People could, if they so choose, adopt any number of strategies involving nearly any number of steps (e.g., two such examples are shown in Fig. 3A-B).

However, specifying these foot placements alone does not capture *how* any given biped might perform these steps. We must also consider the distribution of steps (i.e., *δ_zB_/δ_w_*) at each intermediate step as the stepping goals change (i.e., Δ*z_B_*\* and/or Δ*w*\*). Here, our theoretical framework thus allows us to posit different empirically-testable hypotheses (e.g., Fig. 4D-F) about how these *δ_zB_/δ_w_* ratios should also change at each step. For example, a walker that strongly prioritized regulating step width (e.g., Fig. 4A), as humans do in straight-ahead walking [1], could maintain that strategy but then simply take first one step and then a second, each with an appropriately larger step width (Fig. 4D). Alternatively, the walker could instead seek to simply take the first step to the new stepping goal for that foot, followed by an appropriate second step (Fig. 4E). If on the other hand, a walker wanted to trade-off stability to maximize maneuverability [8, 9], foot placement at the intermediate step should be as accurate as possible to minimize stepping errors at the final step. Theoretically then, stepping distributions at this ideal intermediate step should be approximately *isotropic* (i.e., circular), reflecting no preference for either step width or position regulation (Fig. 4F). One could imagine similarly proposing other alternative competing hypotheses.

### Testing Theoretical Predictions in Humans

In our lane-change experiment, we observed a four-step strategy to be most typical, where participants took a large primary transition step and smaller preparatory and recovery steps (Fig. 2). Incorporating the ideas of the previous section, we can then make theoretical predictions (e.g., as in Fig. 4D-F) about what such a stepping strategy might look like in the [*z_L_, z_R_*] plane. We can then test those predictions against our data.

First, given a reasonable estimated lateral offset (ε) for the preparatory and recovery steps (see Methods), we can determine the foot placements, [*z_L_, z_R_*], and corresponding transition stepping goals, [*z_B_*, w*\*], at each intermediate step (Fig. 5A). Second, we hypothesize that while humans strongly prioritize maintaining step width to maintain stability, [1], they would also trade-off that stability to maximize maneuverability [8, 9] to execute this lateral lane change maneuver. Because they took more than the minimum 2 steps to execute the maneuver, we do not expect the intermediate steps to exhibit perfectly isotropic idealized distributions (e.g., as in Fig. 4F). Taking more intermediate steps during a maneuver reduces the accuracy required at each individual step because those errors can be corrected at subsequent intermediate step(s) prior to reaching the final goal. Hence, we would thus predict these stepping distributions should be *most* isotropic at the larger primary transition step, but intermediately isotropic at the smaller preparatory and recovery steps (Fig. 5A). We thus hypothesized that humans would regulate their stepping movements across these lane-change maneuvers in accordance with these theoretical predictions.

**Figure 5.**
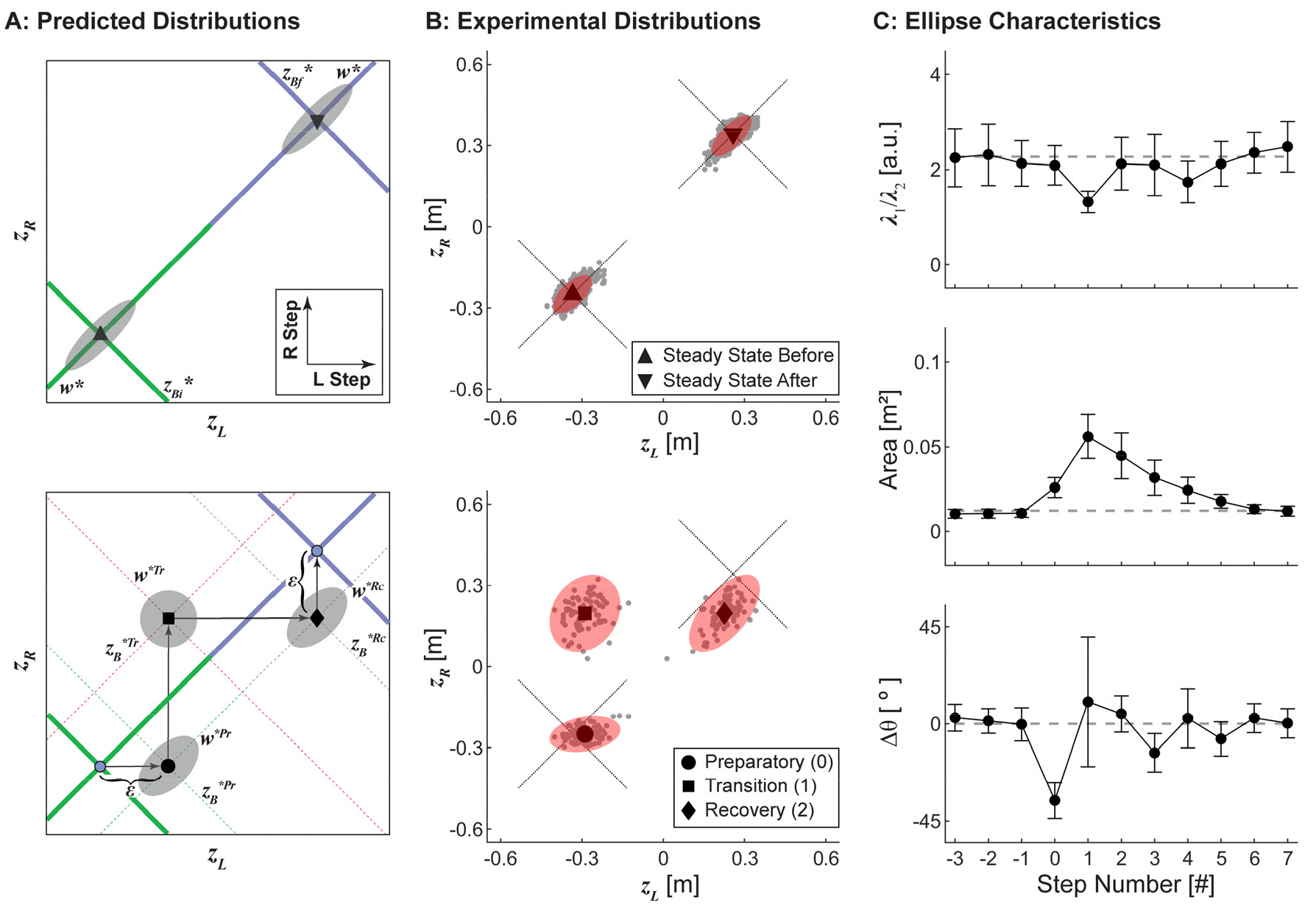
Experimental Stepping Distributions. A) Predicted stepping goals and distributions for the experimentally-imposed lane-change maneuver during both the steady-state walking periods (top) and transition steps (bottom). During steady-state walking before and after the maneuver, we predicted stepping distributions would be strongly aligned to the *w*\* GEM, reflecting strong prioritization of step width over position regulation, as we have previously observed [1]. Participants most commonly performed the maneuver using a four-step maneuver strategy (Fig. 2), which included 3 intermediate steps with distinct stepping goals. Stepping distributions were predicted to be most isotropic at the large primary transition step, and intermediately isotropic at the smaller preparatory and recovery steps. B) Experimental stepping data during the steady state periods before and after the transition (top) and during the preparatory, transition, and recovery steps from 78 analyzed maneuvers, plotted to appear from left to right (bottom), projected onto the [*z_L_, z_R_*] plane. The steady-state data were pooled across 16 steady-state steps before and after the transition from 20 participants performing 4 transitions each (1280 steps total). The diagonal dotted lines indicate the initial and final constant-*z_B_*\* and constant-*w*\* GEMs. The gray dots indicate individual steps, and the red ellipses depict 95% prediction ellipses derived using a *χ*^2^ distribution. C) Experimental 95% prediction ellipse characteristics at each step: aspect ratio (top), calculated as the ratio of the eigenvalues of the covariance matrix of the ellipse, area (center), and orientation (bottom), calculated as the angular deviation (positive angles indicate counter-clockwise) from the orientation of the constant-*w*\* GEM. Error bars indicate ±95% confidence intervals at each step derived using bootstrapping. Gray dashed lines indicate the mean of each characteristic from the two steady state ellipses in (B; top).

We tested this hypothesis by plotting experimental data in the [*z_L_, z_R_*] plane and fitting 95% prediction ellipses to the data for each relevant step (Fig. 5B). During steady-state walking both before and after the maneuver, participants’ steps were strongly aligned to the constant-*w*\* GEM (Fig. 5B; top), consistent with the expected strong prioritization of step width regulation [1]. During each of the preparatory, transition, and recovery steps, the locations of the experimental stepping distributions were consistent with the intersections of the predicted *z_B_*\* and *w*\* GEMs at each step (Fig. 5B; bottom). Furthermore, the experimental stepping distributions themselves (Fig. 5C) were qualitatively consistent with our theoretical predictions (Fig. 5A). The fitted 95% prediction ellipses at the preparatory and transition steps were significantly more isotropic than that observed during steady-state walking (Fig. 5C; top), with the primary transition step being most isotropic as theoretically predicted. Thus, these experimental data strongly support our hypothesis that humans regulate their movements from one step to the next during these lane-change maneuvers by adapting their stepping goals and the relative weighting of *z_B_* and *w* regulation.

### Modeling Adaptive Stepping Regulation for Lateral Maneuvers

Our theoretical predictions (Fig. 4) and confirmatory experimental results (Fig. 5) both strongly suggest that humans can accomplish a range of substantially *non*-steady-state lateral maneuvers by simply changing how they weigh different task-level costs from each step to the next. One does not need to infer some entirely different stepping regulation process to explain the observed behavior. We therefore sought to determine the extent to which our previous multi-objective model [1] could emulate such variations in stepping dynamics if we allowed the parameters of our model to adapt from step-to-step.

Our regulator models are linear update equations (see Methods). Their behavior is controlled primarily by three basic parameters: a pair of target values or stepping goals, [*z_B_*, w*\*], a weight that indicates each regulator’s relative importance, *ρ*, and the additive noise, *σ_a_*, that represents the strength of physiological perceptual/motor noise. Here, we conducted three sequential numerical experiments to assess the effects of adapting each of these model parameters on key stepping dynamics: time series, errors, and variance distributions. We first incorporated adaptive stepping goals that we derived theoretically to approximate an idealized four-step transition strategy (Fig. 5A). Next, we adapted the proportionality parameter, *ρ*, to reflect the predicted stepping distributions (Fig. 5A). Finally, we increased the additive noise to reflect the observed increase in variability during the lateral maneuver (Fig. 5B-C). Importantly, we did not attempt to precisely estimate the model parameters, but rather selected parameters based on the idealized theoretical considerations described above. This approach is analogous to using “templates” of legged locomotion to reveal the basic principles of human walking by testing fundamental hypotheses about the underlying regulation strategies [53, 54].

We algebraically derived (see Methods) new stepping goals, [*z_B_*, w*\*], for each consecutive step (Fig. 6A) that approximated an idealized four-step lane-change maneuver strategy (Fig. 5A). These derived stepping goals included a slightly narrower *w*\* and slight shift in *z_B_*\* towards the new path at the preparatory step, a much wider *w*\* and large shift in *z_B_*\* towards the new path at the transition step, and a slightly narrower *w*\* and slight shift in *z_B_*\* towards the new path again at the recovery step. All other model parameters were assigned constant values across all steps (Fig. 6A), equivalent to the original stepping regulation model [1]. Likewise, during the steady-state periods before and after the lateral maneuver, all model parameters were also held constant to reflect steady-state walking [1].

**Figure 6.**
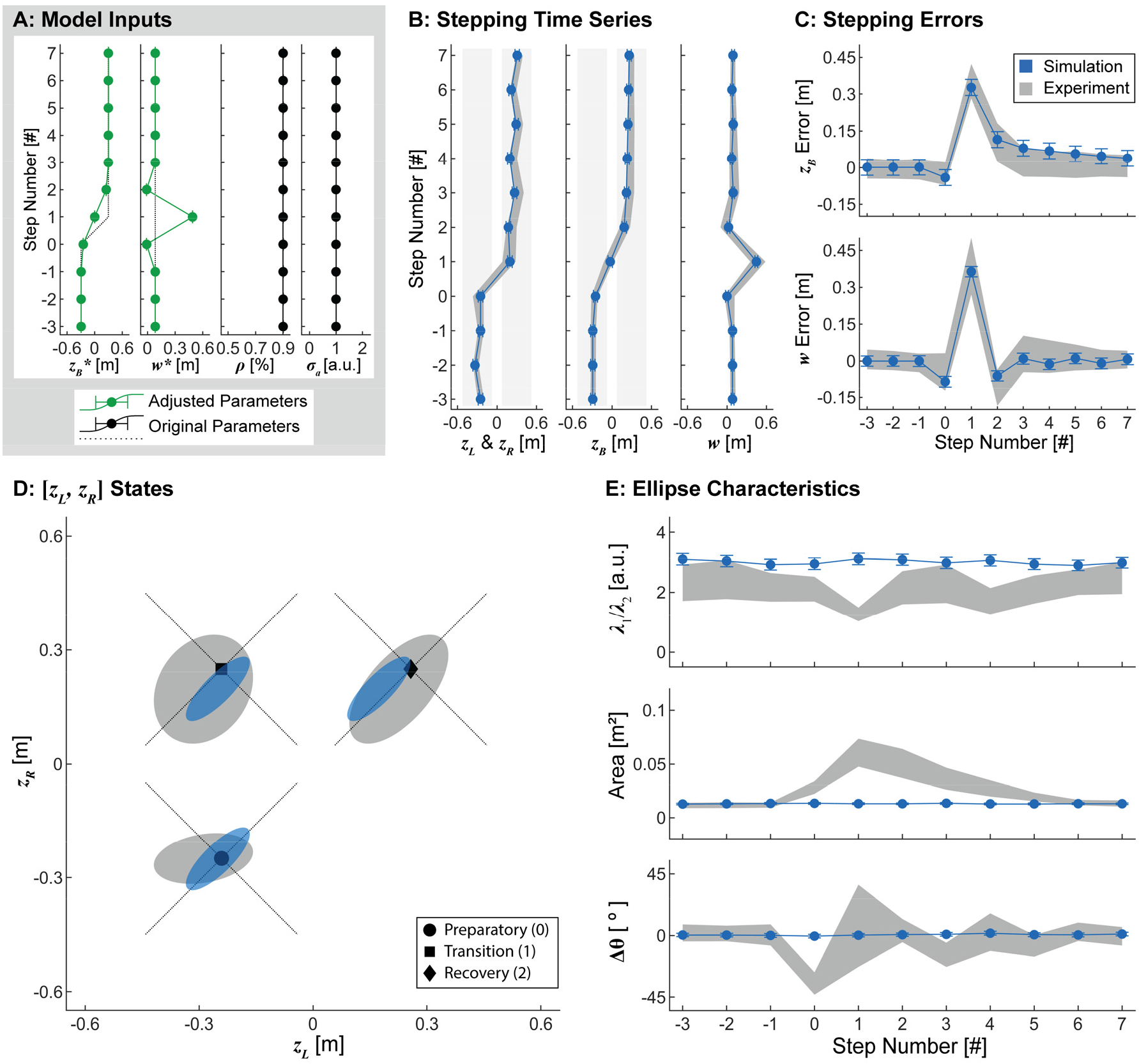
Adaptive Stepping Goals. A) Model parameters. Here, stepping goals, [*z_B_*\*, *w*\*] were updated at each step to reflect an idealized four-step transition strategy. The control proportion (*ρ*) and additive noise (*σ_a_*) were held constant across all steps. B) Stepping time series (mean ± SD) of 1000 simulated lateral transitions using the parameters in (A). C) Stepping errors (mean ± SD) for the simulations in (B). In both (B) and (C), gray bands indicate the middle 90% range from experimental data. D) Simulated stepping data from the preparatory, transition, and recovery steps projected onto the [*z_L_,z_R_*] plane. Gray ellipses represent 95% prediction ellipses at each step from the experimental data. Blue ellipses represent 95% prediction ellipses from the 1000 simulated lateral transitions. The diagonal dotted lines indicate the predicted constant-*z_B_*\* and constant-*w*\* GEMs at the preparatory, transition, and recovery steps. E) Ellipse characteristics (mean ± SD) at each step (as defined in Fig. 5): aspect ratio (top), area (center), and orientation (bottom). Gray bands indicate ±95% confidence intervals from the experimental data derived using bootstrapping. Adapting the stepping goals alone yielded experimentally plausible stepping time series (B), errors (C), and locations (D), but not stepping distributions (D). Thus, adaptive stepping goals are necessary but not sufficient to replicate human stepping during lateral maneuvers.

As expected, allowing the stepping goals to adapt at each consecutive step emulated the experimentally observed stepping time series and errors, as well as the locations of the stepping distributions at each step of the lateral maneuver (Fig. 6B-D). These adaptations captured the proposed feasible strategy of maintaining strong *w*\* prioritization, while taking 2 consecutive steps with appropriately larger step widths (Fig. 4D). However, these parameter variations did not influence the shape, area, or orientation of the stepping distributions, features we did observe experimentally (Fig. 6D-E). Thus, allowing the stepping *goals* to adapt from step to step appears necessary, but is not sufficient to elicit experimentally plausible stepping dynamics during lateral maneuvers.

Next, we added the ability to adaptively modulate *ρ* at each step (Fig. 7A) to incorporate our theoretical predictions for an idealized four-step lane-change maneuver (Fig. 5A). Typical steady-state walking heavily weights regulating step width over lateral position (i.e., *ρ* ≈ 0.9) [1], presumably to maintain lateral stability. Conversely, we expect people to trade off stability to gain maneuverability [8, 9] while executing this maneuver. We therefore set *ρ* = 0.50 at the transition step (Fig. 7A) to specify equal weighting of step width and position regulation (Fig. 4F), thereby maximizing maneuverability. We then set *ρ* = 0.7 at the preparatory and recovery steps (Fig. 7A) to reflect an intermediate multi-objective cost weighting (Fig. 5A). The same adaptive stepping goals (Fig. 6) were again incorporated here. All other model parameters were assigned constant values across all steps (Fig. 7A).

**Figure 7.**
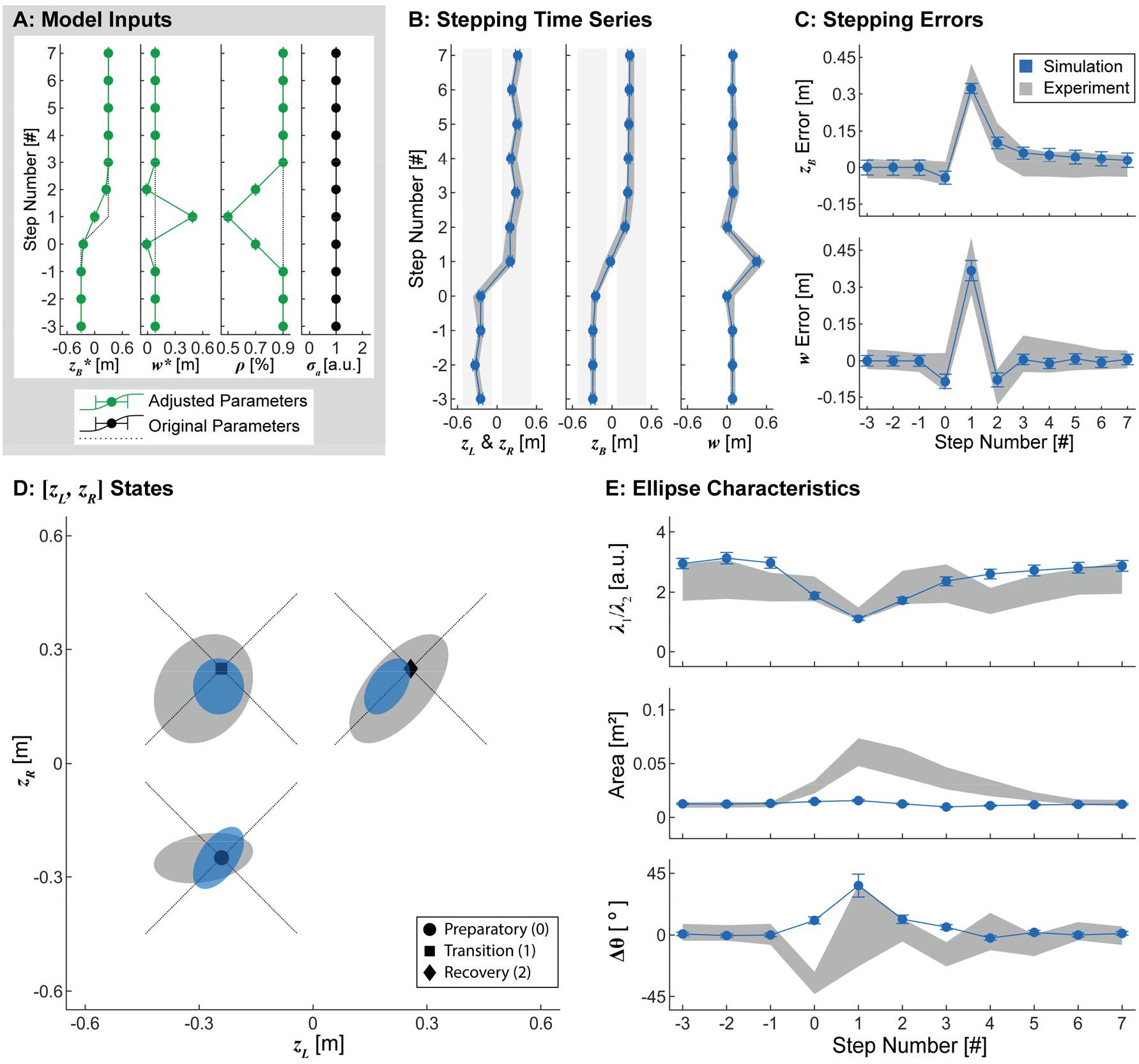
Adaptive Control Proportion. A) Model parameters. Here, in addition to adapting the stepping goals ([*z_B_*\*, *w*\*]; Fig. 6), the control proportion (*ρ*) was also varied during the preparatory, transition, and recovery steps to reflect the hypothesized maneuverability and error correction at each step. Additive noise (*σ_a_*) remained unchanged from the original, constant parameter model. B-E) Results obtained from 1000 simulated lateral transitions using the parameters in (A), with data plotted in an identical manner to Fig. 6B-E. Adding step-to-step modulation of ρ again captured experimental stepping time series and errors at each step (B-C). Here however, allowing *ρ* to adapt at each step also induced changes in prediction ellipse aspect ratios during the transition steps that were qualitatively similar to those observed experimentally. Modulating *ρ* however, did not induce corresponding changes in ellipse area (D-E). Thus, adapting *ρ* at each step is also necessary but not sufficient to emulate human stepping during lateral maneuvers.

Allowing both the stepping goals and *ρ* to adapt at each consecutive step again captured the experimentally observed stepping time series, stepping errors, and stepping distribution locations at each step of the lateral maneuver (Fig. 7B-D). Furthermore, modulating *ρ* produced more isotropic simulated stepping distributions at the preparatory, transition, and recovery steps, consistent with both our theoretical expectations (Fig. 5A) and experimental observations (Fig. 5B-C). However, these simulations did not influence the area of the stepping distributions (Fig. 7D-E), and the changes to orientation were not consistent with what was observed experimentally (Fig. 5C). Thus, in addition to adapting stepping goals (Fig. 6), adapting *ρ* from step to step is also necessary, but still not yet sufficient to emulate human stepping dynamics during this lateral maneuver.

Therefore, in addition to adapting both the stepping goals (Fig. 6) and *ρ* (Fig. 7) from step to step, we then doubled the additive noise (*σ_α_*) in the model at the preparatory and transition steps (Fig. 8A). Additive noise is thought to reflect physiologic noise from a variety of sources, including sensory, perceptual, and/or motor processes [30]. Here, we assessed whether increasing this additive noise could emulate the increases in the stepping distribution areas observed experimentally (Fig. 5B-C).

**Figure 8.**
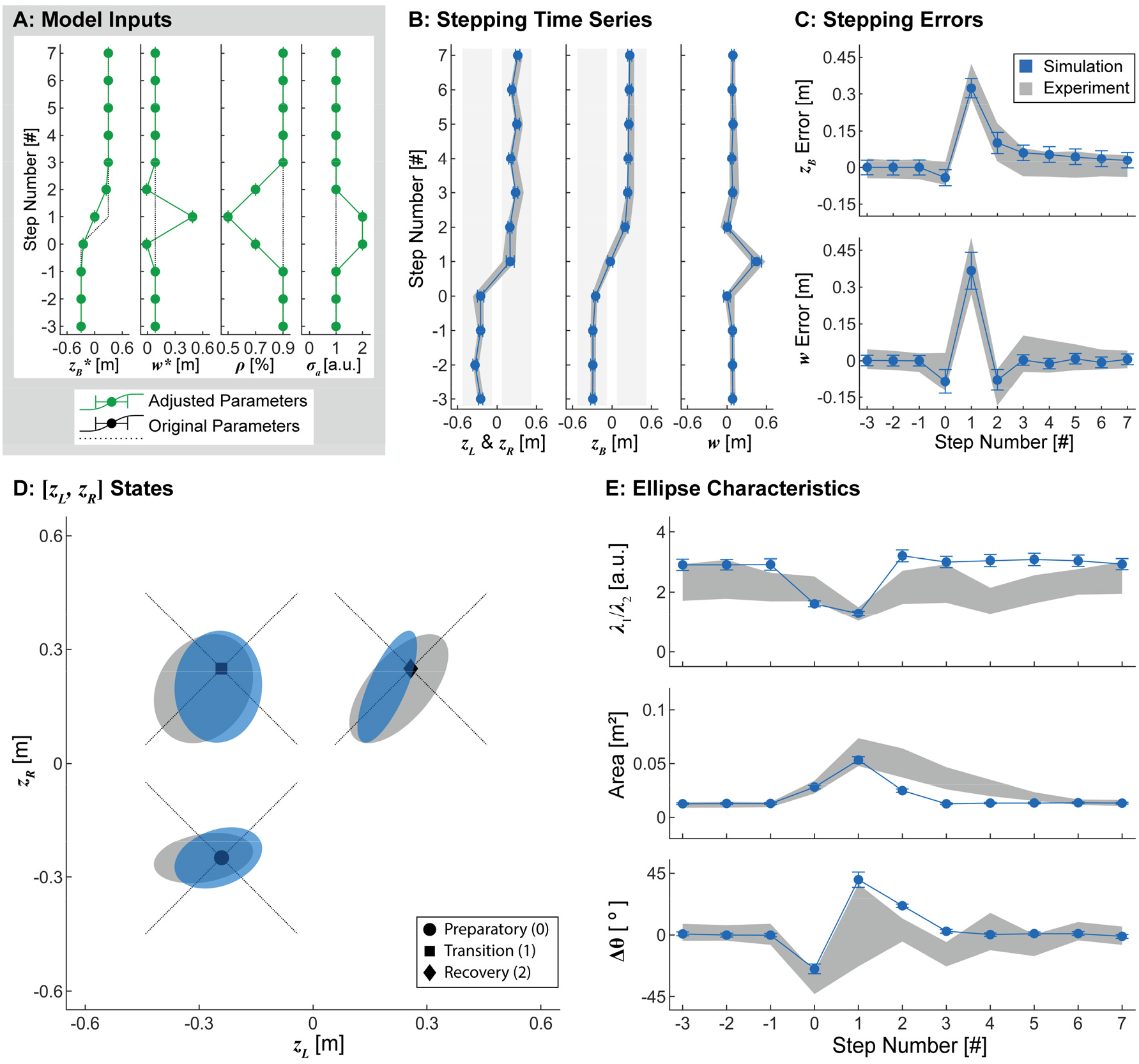
Adaptive Additive Noise. A) Model parameters. Here, in addition to adapting the stepping goals (Fig. 6) and *ρ* (Fig. 7), additive noise (*σ_a_*) was also doubled at the preparatory and transition steps. B-E) Results obtained from 1000 simulated lateral transitions using the parameters in (A), with data plotted in an identical manner to Figs. 6-7. Adding *σ_a_* modulation again emulated experimental stepping time series and errors (B-C), as well as the qualitative changes in the aspect ratio of the fitted ellipses during the maneuver (D-E). Furthermore, modulating *σ_a_* emulated the experimentally observed increases in ellipse areas. Modulating *σ_a_* also affected orientations of the fitted ellipses, although not entirely in the same ways as the experimental data (D-E). Therefore, adaptively modulating the stepping goals ([*z_B_*\*, *w*\*]), control proportion (*ρ*), and additive noise (*σ_a_*) in our existing model can elicit changes in stepping dynamics qualitatively similar to those observed in humans during this lateral maneuver.

Allowing the stepping goals, *ρ*, and *σ_a_* to adapt at each consecutive step again captured the experimentally observed stepping time series, stepping errors, and stepping distribution locations at each step of the lateral maneuver (Fig. 8B-D) and qualitatively captured the shapes of the stepping distributions (Fig. 8D-E). Increasing *σ_a_* also increased the areas of the stepping distributions at the preparatory, transition, and recovery steps (Fig. 8D-E). Interestingly, increasing *σ_a_* also influenced the orientations of these distributions, inducing a clockwise rotation at the preparatory step and a counterclockwise rotation at the transition step (Fig. 8D-E). These rotations reflect the individual steps at which the increased variability was applied. Increasing the noise at the preparatory step (shown here as taken with the left foot) can only increase variability in the horizontal direction in the [*z_L_, z_R_*] plane. Conversely, increasing the noise at the subsequent transition step (taken with the right foot) can only increase variability in the vertical direction.

We designed these simulations using the basic concepts from our theoretical framework (Fig. 4) to pose specific predictions for an idealized four-step lane-change maneuver (Figs. 5A), comparable to what we observed in our experiment (Fig. 2). Taken together, the results demonstrate that adapting the stepping goals ([*z_B_*, w*\*]; Fig. 6), the relative weight given to regulation for of those goals (*ρ*; Fig. 7), and the additive noise (*σ_a_*; Fig. 8) allowed our simulations to emulate the key changes in stepping dynamics observed during this lateral maneuver task (Fig. 5).

## DISCUSSION

Understanding how humans perform accurate, goal-directed walking movements in the face of inherent variability, redundancy, and equifinality remains a fundamental question in human motor neuroscience. In pursuit of this aim, we previously developed a theoretical framework to describe how humans regulate stepping to achieve continuous, straight-ahead walking [1]. Models developed from this framework provide goal-directed “stepping regulation templates” that are both analogous and complimentary to mechanical templates that describe the *within*-step mechanics and dynamics of walking (e.g., [21, 32, 33]). Here, because humans rarely perform long bouts of steady-state walking [47, 48], we therefore examined if our basic regulation template scheme could be used to model human stepping dynamics during a prescribed lateral maneuver that, by necessity, strongly deviated from steady state motion.

Our basic stepping regulation framework has demonstrated that simple low-dimensional, linear, single-step regulators adequately capture straight-ahead steady-state walking dynamics [1, 30]. Conversely, walking maneuvers involve substantial *non*-steady-state movements that would seem to violate these key assumptions. This could well necessitate the need for higher-dimensional models, nonlinear responses, and/or responses that act across multiple steps to enact any such maneuver. For humans, it is then certainly plausible they might use one control strategy to regulate steady-state stepping (e.g., [1, 30]), switch to some completely different strategy to execute a particular lateral maneuver, then switch back again to their previous strategy to return to steady-state. Here, however, we demonstrate to the contrary that this is not required in this context.

Our *key theoretical contribution* is that we show how our previously developed theoretical stochastic optimal control framework can be modified to model how humans adapt their lateral stepping to enact non-steady-state lateral maneuvers (Fig. 4). Specifically, we show this can be accomplished by adding an additional layer to our previous control hierarchy: namely, the processes responsible for adapting, on a step-by-step basis, the cost landscapes governing stepping regulation. Our *key empirical contribution* is we then demonstrate (Fig. 5) that humans do indeed execute lateral maneuvers in a manner consistent with our theoretical predictions. Our *key computational contribution* is that we show that allowing the parameters in our lateral stepping regulation model to adapt from each step to the next can emulate the changes in lateral stepping dynamics exhibited by humans (Figs. 6-8). To our knowledge, our results are the first to demonstrate that humans might use evolving cost landscapes in real time to perform such an adaptive motor task and, furthermore, that such adaptation can occur quickly – over only one step.

In our simulations (Figs. 6-8), we chose model parameters based on idealized theoretical considerations (Fig. 4), only loosely related to general trends observed in our experiment (Fig. 5A). While our resulting simulations did not precisely “fit” the experimental data, they were not intended to. On the contrary, our aim was to demonstrate that reasonable values of the parameters could yield the same basic dynamical and statistical stepping *features* that humans exhibited during this lateral maneuver task (Fig. 8). This approach is directly analogous to using mechanical “templates” of within-step dynamics of legged locomotion to reveal basic principles of walking [53, 54] and to propose fundamental hypotheses about what high-level control strategies might be acting. Here, we used our theoretical framework to derive such hypotheses (Fig. 4). We then used both experiments (Fig. 5) and simulations (Figs. 6-8) to validate the theoretical predictions facilitated by our templates [1] for an entirely new locomotor context.

Many studies have used optimal control theory to model a myriad of individual motor tasks (e.g., [52, 55, 56]). Such efforts, however, did not consider tasks where the task goals and/or movement objectives change as the task is being performed. Likewise, many studies have addressed motor adaptation (e.g., [57, 58]) and/or motor learning (e.g., [59, 60]), including for walking tasks (e.g., [61–63]). These paradigms, however, track how task performance changes slowly over many repetitions (typically at least 10’s to 100’s), and not from one repetition to the next. Furthermore, while multiple studies have demonstrated that such longer-term adaptation can be replicated by “lag-1” type computational models that explicitly correct only for errors experienced on one previous iteration of the task (e.g., [64–66], those models were not *themselves* “adaptive”. On the contrary, they presumed some constant process (with constant model parameters) that achieved adaptation over multiple repetitions of the task under consideration. In sum, none of these extremely well-trodden paradigms quite capture the nature of the task we studied here.

Conversely, rapid adaptation in response to changing environmental contexts has been demonstrated in both birdsong [67–69] and human speech [70]. This apparent plasticity of a well learned, crystalized behavior suggests that vocalization may be controlled by a “malleable template,” in which trial-by-trial variability is used to adapt learned behaviors [71]. Such rapid adaptations have also been observed in both animal [15–17] and human [7–9] locomotion (i.e., maneuvers). As the associated motor planning processes occur nearly instantaneously [72–74], this prior work supports our findings that humans can and do make rapid adaptations to their stepping regulation to enact lateral maneuvers. Thus, in demonstrating that our lateral stepping regulation framework successfully predicts how humans perform lateral maneuvers, our findings support the notion that humans use a malleable template [71] to make “embodied decisions” [6] about how to regulate their stepping movements in real time during ongoing locomotion.

Prior work has postulated the existence of a “stability-maneuverability trade-off” during lateral maneuvers [7, 8]. However, in the absence of a coherent, predictive, theoretical framework, this stability-maneuverability trade-off has not been adequately defined, much less confirmed. We propose that our lateral stepping regulation framework, and the models derived from it, provides the necessary theoretical and computation foundation needed to describe how humans trade-off stability for maneuverability during lateral movements. By extension, our findings suggest that stability and maneuverability in the context of locomotion are not distinct and independent concepts, but rather different manifestations of the same underlying stepping regulation process. Indeed, if stability and maneuverability were independent, as often assumed, humans should be able to remain stable and maneuverable simultaneously. However, our theoretical framework (Fig. 4) demonstrates precisely how and why humans must trade-off some degree of stability (i.e., *w*-regulation) to gain the maneuverability needed to perform lateral maneuvers.

Walking in the real world often requires maneuverability to adapt to changing environmental conditions or goals. We suggest that such maneuvers are governed by a hierarchical control/regulation schema with at least three distinct layers: low-level processes that govern within-step dynamics to ensure viability [34, 36], step-to-step regulation to achieve goal-directed walking [1, 30, 34, 35], and the presently demonstrated modulation of stepping regulation to achieve adaptability. The ability of our lateral stepping regulation framework to emulate human stepping during the lane change maneuver studied here demonstrates that its predictive capabilities extend to a much greater range of walking tasks than initially thought, encompassing not just steady state walking, but transient behaviors as well.

## METHODS

### Ethics Statement

Prior to participating, all participants signed written informed consent statements approved by the Institutional Review Board at The Pennsylvania State University (PSU IRB Study #00011140).

### Lateral Maneuvers in Humans

The data analyzed here were collected as part of a previous experiment involving twenty young healthy adults (Table 1) [51]. All participants were screened to ensure they had no history of orthopedic problems, recent lower extremity injuries, any visible gait anomalies, or were taking medications that may have influenced their gait.

**Table 1.** Participant Characteristics. All values except Sex are given as Mean ± Standard Deviation.

| Characteristic: | Value: |
| --- | --- |
| Sex | 9 M / 11 F |
| Age [yrs] | 21.7 ± 2.6 |
| Body Height [m] | 1.73 ± 0.08 |
| Body Mass [kg] | 69.9 ± 12.1 |
| Body Mass Index (BMI) [kg/m<sup>2</sup>] | 23.2 ± 2.7 |
| Leg Length [m] | 0.82 ± 0.06 |

The experimental protocols were described in detail previously [51]. Briefly, participants walked in an “M-Gait” system, comprised of a 1.2m wide motorized treadmill in a virtual reality environment (Motek, Amsterdam, Netherlands). Each participant walked at a constant speed of 0.75 m/s. Following a 4-minute acclimation trial, participants completed several different walking trials involving path navigation. The data analyzed here were generated from one such trial, during which participants were instructed to switch between two parallel paths, centered 0.6m apart, following an audible cue (Fig. 2). Participants completed 6 maneuvers during one 4-minute walking trial, and were instructed to walk normally on their current path between maneuvers. The first and last maneuvers occurred too close to the beginning and end of the walking trial, respectively, to ensure participants were walking normally both before and after the maneuver. Therefore, the present analysis included only the middle four maneuvers from each participant, for a total of 80 lateral maneuvers.

Kinematic data were recorded from 16 retroreflective markers placed on the head, pelvis, and feet of each participant. Marker trajectories were collected at 120Hz using a 10-camera Vicon motion capture system (Oxford Metrics, Oxford, UK) and post-processed using Vicon Nexus and D-Flow software (Motek, Amsterdam, Netherlands). Marker trajectories were analyzed in MATLAB (MathWorks, Natick, MA). Heel strikes were determined using a velocity-based detection algorithm [75]. Lateral foot placements (*z_L_* and *z_R_*) were defined as the lateral location of the heel marker at each step. Step width (*w*) and lateral position (*z_B_*) were then determined at each step using Eq (1) (Fig. 1C).

We were unable to determine accurate heel strikes for one lateral maneuver and consequently excluded this maneuver from further analyses. Stepping data from the remaining 79 maneuvers were normalized to a constant direction (left-to-right) and to the initiation of the transition, defined here as the last step taken on the original path. Means and standard deviations of step width and position were determined at each step of the lateral maneuver. The steady-state mean step width was determined from all participants and all steps, excluding the 5 steps before to 10 steps after each transition. The steady-state mean position was determined similarly, but was determined separately for each path (left and right). These steady-state means were used to define the stepping goals for this task, [*z_B_*, w*\*]: *w*\* was defined as the steady-state mean step width, and *z_B_*\* before and after the maneuver was defined as the steady-state mean position on the left path and the steady-state mean position on the right path, respectively. Errors in both position and step width (mean±SD) were determined at each step of each maneuver as the differences in each from these steady-state stepping goals (Fig. 2C).

### Model of Lateral Stepping Regulation

To model lateral stepping, the simplest, mechanically sufficient biped includes the lateral locations of the body (*z_Bn_*) and each of the two feet (*z_Ln_* and *z_Rn_*; Fig. 1B) [33, 76, 77]. We presume that left and right foot placement are coordinated to achieve some more general walking task goal or goals. One such goal is to maintain lateral balance by regulating step width (*w_n_*; Fig. 1C) [23, 32, 78, 79]. Humans also regulate the lateral position of the body with respect to their path [20, 77, 80, 81], which can be approximated by the midpoint between the two feet during upright walking [76, 78]. Here, we define the body position for a single step (*z_Bn_*, Fig. 1C) as the midpoint between the two feet [1, 82]. This definition of *z_Bn_* is strongly correlated to the lateral center-of-mass (CoM) position at each heel strike ([83]; Supplement). By selecting appropriate foot placements, {*z_L_, z_R_*}, humans can regulate both their body position and step width,{*z_Bn_, w_n_*} [1], via:

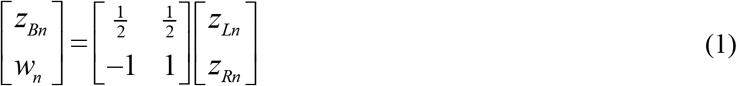

During continuous straight-ahead walking, multi-objective regulation of primarily step width and secondarily lateral position captures human lateral step-to-step dynamics [1]. The regulation model minimizes errors with respect to both step width and lateral position using the goal functions [37, 38, 40]:

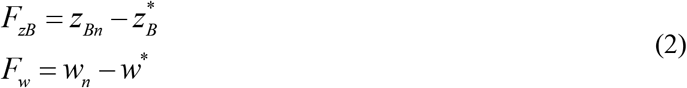

where the goal is to drive each *F* → 0. The value of each regulated variable at the subsequent step is determined independently using the state update equations:

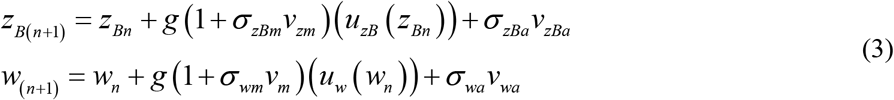

where *σ_mv_m* and *σ_av_a* represent multiplicative and additive noise terms, respectively.

The control inputs, *u_z_B*(*z_Bn_*) and *u_w_*(*w_n_*), were derived analytically as stochastic optimal single-step regulators with direct error feedback [1, 30, 31, 40], following the Minimum Intervention Principle [41, 52]. Such controllers are optimal with respect to the following quadratic cost functions:

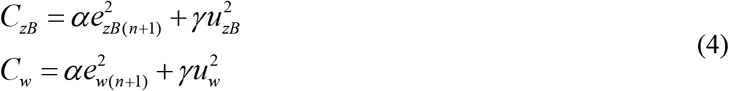

The first term of each cost function penalizes errors with respect to the goal function (Eq (2)) at the next step.

The second term penalizes “effort”, quantified as the magnitude of the control input. Here, *α* and *γ* were positive constants that weighted the two terms of each cost function [1]. The subsequent analytical derivation [1] yielded control inputs as:

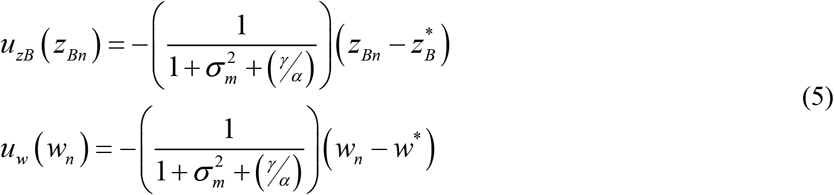

The value of each regulated variable, [*z_B_, w*], at the subsequent step is determined independently using separate, individual optimal controllers. Thus, two corresponding foot placements are determined, one using the predicted value of position, and the other using the predicted value of step width. The final predicted value of foot placement is determined as a weighted average (*ρ*) of the predictions from each regulated variable, where *ρ* = 0 indicates 100% position control and *ρ* = 1 indicates 100% step width control [1]. Hence, stepping regulation is conceived here as arising from a “mixture of experts” [84, 85]. In this way, the value of *ρ* directly determines the shape of the cost landscape, as evidenced by the changing shapes of the variability ellipses in the [*z_L_, z_R_*] plane (Fig. 7).

As in our prior work [1], the following parameters were set to constant values for all simulations. The baseline amplitude of the additive noise (*σ__a_*) was set to the steady-state standard deviation of each variable. The amplitude of the multiplicative noise (*σ__m_*) was set to 10% of the additive noise level (i.e., 0.1·*σ__a_*). The magnitude of the additive and multiplicative noise at a given step were determined from the Gaussian random variables *v__a_* and *v__m_*, respectively, each with zero mean and unit variance. Error correction was weighted much more heavily than effort in the cost function by setting γ/α = 0.10.

### Steady-State Stepping Regulation

We determined if any constant value of *ρ* could emulate human stepping throughout the lateral maneuver (Fig. 3C). For each value of *ρ* between *ρ* = 0.00 and *ρ* = 1.00, in increments of *Δρ* = 0.01, we conducted 1000 batches of 1000 simulations of the lateral maneuver. For each batch, we determined the means and standard deviations of both position and step width at each step. We then computed the overall mean and standard deviation of each variable at each step across all batches. For each variable, simulated and experimental values were compared by plotting the simulated overall mean ± 1SD against 95% confidence intervals of the experimental data determined from bootstrapping [40, 86]. For a given value of *ρ*, we considered the model successful at capturing human stepping if the simulated means and standard deviations of step width and position fell within the corresponding experimental 95% confidence intervals at every step of the lateral maneuver (Fig. 3C).

### Stepping Regulation for Non-Steady-State Tasks

When the *z_B_*\* and *w*\* GEMs are viewed in the [*z_L_, z_R_*] plane, it is clear that at least one intermediate step is necessary to accomplish any Δ*z_B_*\* and/or Δ*w*\* maneuver (Fig. 4A). For the minimum two-step maneuver strategy (Fig. 4B), the stepping goals for either transition step can be determined algebraically from the steady-state initial, [*z_Bi_*\*, *w_i_*\*], and final, [*z_Bf_*\*, *w_f_*\*], stepping goals. We first determined the initial and final foot placement goals from the initial and final stepping goals using Eq (1). The foot used to take the intermediate step must be placed at that foot’s final foot placement goal, while the stance foot remains at its initial foot placement goal. For example, if the right foot is used to take the intermediate step (Fig. 4B; a), the intermediate foot placement goals are [*[z_Li_*\*, *z_R_*\*]. By transforming these intermediate foot placement goals back into [*z_B_, w*] coordinates using Eq (1), and re-defining the final stepping goals as the initial stepping goals plus the changes in the stepping goals, [Δ*z_B_*\*, <u>Δ</u>*w*\*], the intermediate stepping goals are:

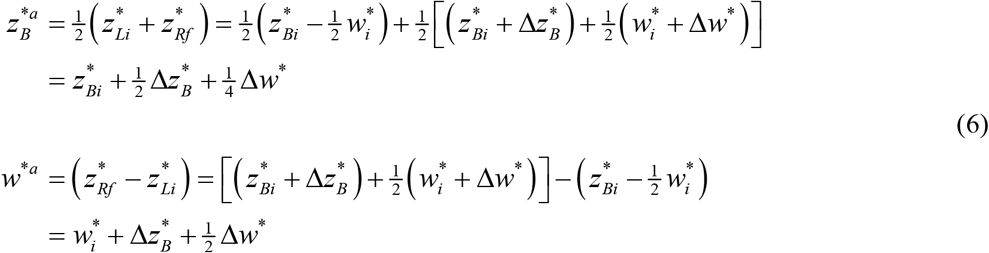

The same procedure can be used to determine the stepping goals for an intermediate step taken with the left foot (Fig. 4B; b) by simply reversing the initial and final components of the intermediate foot placement goals and making the corresponding substitutions in Eq (6).

However, equifinality exists in the number of steps and their placements that can be used to accomplish any Δ*z_B_*\* and/or Δ*w*\* maneuver. Experimentally, we most commonly observed a four-step maneuver strategy with distinct preparatory, transition, and recovery steps (Figs. 2, 4C). To determine the stepping goals associated with an idealized version of such a four-step strategy, we first calculated a single intermediate offset (*ε*) by averaging the experimental differences in foot placement at each preparatory and recovery step relative to steady-state walking. Assuming a left-to-right transition with preparatory and recovery steps that were each taken with the left foot, the foot placement goals at the preparatory, [*z_L_^*Pr^, Z_R_^*Pr^*], and recovery, [*z_L_^*Rc^, z_R_^*Rc^*], steps were defined as [*z_Li_* +*ε*, z_Ri_*\*] and [*z_Lf_*, Z_Rf_*\*-*ε*], respectively. These foot placement goals were then transformed back into [*z_B_, w*] coordinates using Eq (1) to determine the preparatory, [*z_B_^*Pr^, w^*Pr^*], and recovery, [*z_B_^*Rc^, w^*Rc^*], stepping goals. The transition stepping goals, [*z_B_^*Tr^, w^*Tr^*], were then determined using Eq (6) above, substituting the preparatory and recovery stepping goals for the initial and final stepping goals.

The stepping goals themselves, however, do not provide insight into *how* humans regulated their stepping during the lateral maneuver. For a four-step maneuver strategy with a large primary transition step and smaller preparatory and recovery steps, as was observed experimentally (Fig. 2), stepping distributions are theoretically predicted to be most isotropic at the primary transition step and intermediately isotropic at the preparatory and recovery steps.

We qualitatively assessed the accuracy of the theoretically predicted stepping goals and distributions by comparing them to the experimental data at each step, plotted in the [*z_L_, z_R_*] plane (Fig. 5B). The maneuver completed with a large cross-over step was excluded from analyses of the stepping distributions, as this maneuver was completed with a different stepping strategy. Using the remaining 78 maneuvers, we first computed the covariance matrix of left and right foot placements at each step: **C** = cov(***z**_L_, **Z**_R_*). We take {*λ*_1_, *λ*_2_} and {**e**_1_, **e**_2_} to denote the eigenvalues and eigenvectors of **C**, respectively. We then construct a 95% prediction ellipse by scaling the eigenvalues by the 95^th^ percentile critical value of the Chi-Squared distribution:

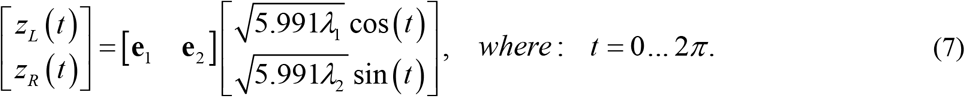

Thus, we expect 95% of the data points to lie inside this ellipse, assuming a bivariate normal distribution. We then characterized each such ellipse by its shape, size, and orientation (Fig. 5C). We defined the shapes of each ellipse by their aspect ratio, computed as the ratio of the major- and minor-axis eigenvalues: *λ*_1_/*λ*_2_. We computed the sizes of each ellipse as their area:

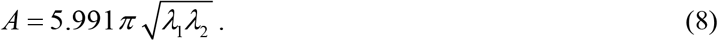

We computed the orientations of each ellipse as the angle of the major axis measured counterclockwise from the *w*\* GEM:

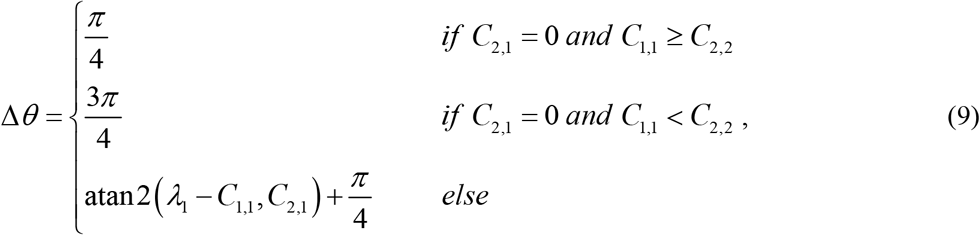

where the *C*_i,j_ are the respective elements of the covariance matrix. For each ellipse characteristic computed at each step, we then calculated 95% confidence intervals for these values using bootstrapping.

### Adaptive Stepping Regulation for Lateral Maneuvers

We adapted parameters of our goal-directed multi-objective lateral stepping regulation model from step-to-step to determine if such adaptions could model the observed stepping dynamics during the lateral maneuver task. For each model iteration, all parameters not explicitly modulated were assigned a constant value in accordance with our previous steady-state stepping regulation model [1]. We first replaced the stepping goals *z_B_*\* and *w*\* in the model (Eq (2)) with the estimated adaptive stepping goals (see previous section) (Fig. 6A). We next adapted *ρ*, the model parameter that specifies the relative weighting of step width and position regulation. The values assigned to *ρ* were chosen based upon theoretical predictions of the maneuverability and error correction requirements at each intermediate step (see previous section; Fig. 5). Specifically, we set *ρ* = 0.50 at the transition step, specifying equal weighting of step width and position regulation. We set *ρ* = 0.7 at the preparatory and recovery steps, a value intermediate between the approximate value observed for steady-state walking (*ρ* ≈ 0.9) and the value of 0.5 used for the transition step (Fig. 7A).

Finally, in addition to the adaptive stepping goals and *ρ* modulation, we doubled the additive noise parameter (*σ_a_*) at the preparatory and transition steps (Fig. 8A). The increased area of the stepping distributions at the preparatory, transition, and recovery steps reflects greater overall variability with respect to both the *w*\* and *z_B_*\* GEMs. Additive noise is thought to reflect physiologic noise from sensory, perceptual, and/or motor processes [30], which is expected to increase during the lateral maneuver task. Here, we increased the additive noise (Fig. 8A) to determine the extent to which this would qualitatively capture the types of increases in stepping distribution areas that we observed experimentally.

For each model iteration, we assessed the ability of the model to emulate the key changes in stepping dynamics observed experimentally: stepping time series (Figs. 6–8B), stepping errors (Figs. 6–8C), and stepping distributions (Figs. 6–8D-E) at each step of the lateral maneuver. The lateral maneuver was simulated 1000 times for each model iteration. All maneuvers were oriented from left-to-right, and the transition step was specified to be taken with the ipsilateral (i.e., right) foot relative to the direction of the transition, consistent with all but one of the experimentally observed maneuvers. Time series (mean±SD) of foot placement (*z_L_* and *z_R_*), position (*z_B_*), and step width (*w*) were calculated at each step of the simulated maneuvers. Errors with respect to both position and step width (mean±SD) were calculated as the difference in the simulated position and step width relative to the stepping goals at each step of the simulated maneuvers. The simulated stepping time series and errors were compared to the middle 90% range of the experimental data at each step. The simulated stepping distributions were characterized by the aspect ratio, area, and orientation of a fitted 95% prediction ellipse (see previous section). Error bars at each step for each variable were calculated as 95% confidence intervals using bootstrapping. The simulated ellipse characteristics were compared to the bootstrapped 95% confidence intervals from the experimental data at each step for each variable.

### Statistical Comparisons

As the experimental and simulated data were structured in very different ways, standard inferential statistics (e.g., t-test, ANOVA, etc.) would not be appropriate to compare these results. Furthermore, we could generate a sufficiently large number of model simulations to ensure small p-values for almost any comparison, thereby diminishing the comparative power of any such assessments. Instead, we used descriptive statistics (e.g., standard deviations, confidence intervals, etc.) to quantify the experimental observation values. We then compared model predictions to these observations. We inferred that model predictions that fell within experimentally observed standard deviations/confidence intervals were statistically consistent with the experimental results. Additionally, the aim of this analysis was to determine if hierarchical adaption of our goal-directed multi-objective lateral stepping regulation models could qualitatively emulate the same *types* of changes in stepping dynamics observed during the lateral maneuver task. Descriptive statistics were sufficient to accomplish this aim.

## DATA

All relevant experimental data, and modeling and analysis codes, are available from Dryad (https://doi.org/10.5061/dryad.tx95x6b1x) [87].

## AUTHOR CONTRIBUTIONS

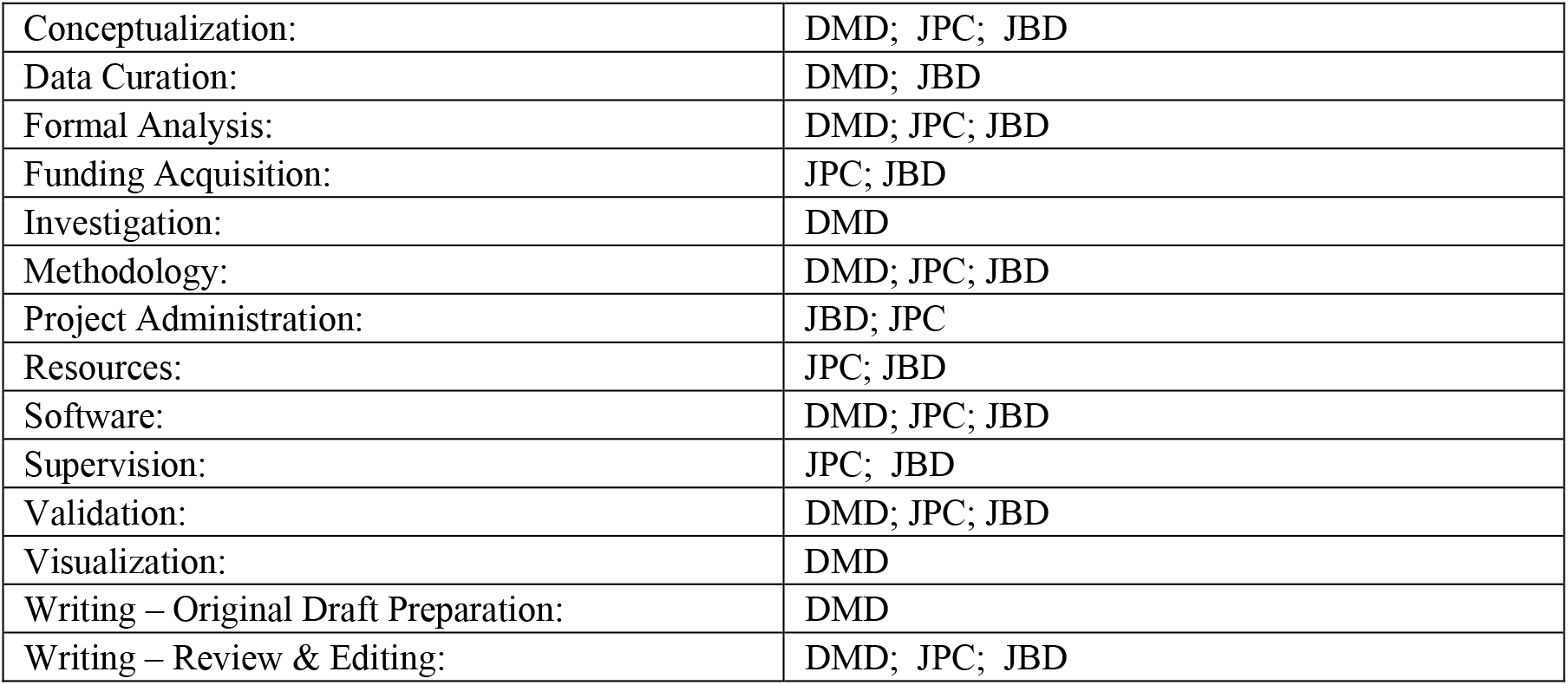

## ACKNOWLEDGEMENTS

The authors thank Dr. Meghan Kazanski and Ms. Anna Render for their assistance with data collection. The authors thank Dr. Meghan Kazanski for her assistance with data processing and initial analyses.

