## Supplement S1 for "Adaptive Multi-Objective Control Explains How Humans Make Lateral Maneuvers While Walking"

***PLoS Computational Biology***

### SUPPLEMENTARY TEXT #S1: Relation of Body Position ($z_B$) to Center-of-Mass (CoM)

We define lateral body position ($z_B$) at each step as the midpoint between consecutive heel strikes ([1]; Fig. 1C). In continuous time, the center-of-mass (CoM) oscillates between the feet as each new step is taken [2, 3]. Near the moment of heel strike, the CoM passes approximately across the midline between the feet, during both straight walking ([3, 4]; Fig. S1A) and when performing lateral maneuvers [5]. In our work, we analyze walking *step-to-step*: each step is treated as a single event. Fig. S1A shows, however, that no single value of CoM value can easily be assigned to a complete step. However, we can take $z_B$ as a *proxy* of a discrete (once per step) estimate of the lateral CoM location for each step (Fig. 1B).

To see this, we computed continuous-time lateral CoM trajectories from pelvis centroid motion [6] (Fig. S1B). We then correlated the lateral CoM values that occurred at each heel strike with $z_B$ for that step (Fig. S1C). In all cases, $z_B$ was very strongly correlated with these CoM values, for steady straight-ahead walking ($R^2 = 0.82$) and also for each of the preparatory ($R^2 = 0.81$), transition ($R^2 = 0.76$), and recovery ($R^2 = 0.82$) steps taken during the lane-change maneuver.

These data confirm that $z_B$ is a reasonable proxy that represents a discrete (i.e., once per step) measure of the approximate lateral position of the body's center-of-mass (CoM) at each step.

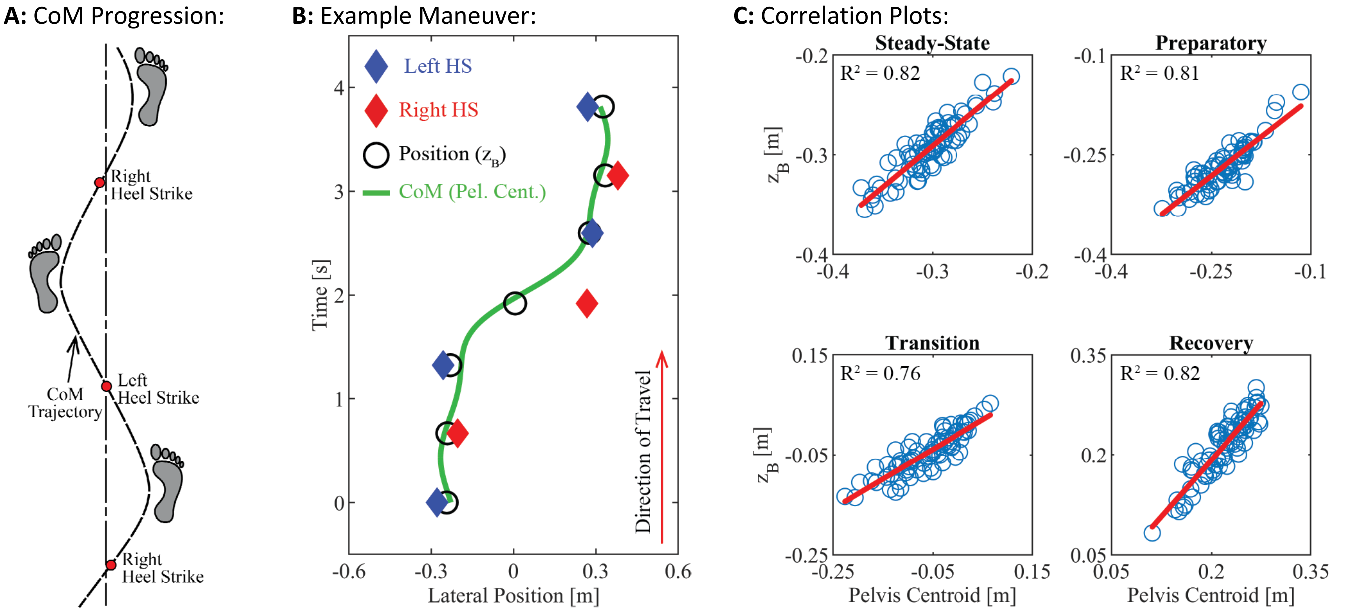


**Figure S1: A:** Schematic of foot placements and center-of-mass (CoM) trajectory during typical straight-ahead walking, with instances of left and right heel strikes indicated (see also [3], Fig. 15 and [4], Fig. 2). **B:** Data for Left (♦) & Right (♦) steps, lateral position ($z_B$) locations (○), and CoM (pelvis centroid) trajectory (—) for a typical lane-change trial from this experiment (similar trends shown in [5], Fig. 2). **C:** Correlations of body CoM at each heel strike vs. $z_B$. Note: All $R^2 \geq 0.76$.
